# Estimating chronological and brain age using risk-taking behavior under uncertainty

**DOI:** 10.64898/2026.03.12.711461

**Authors:** Yunchen Gong, Meiling Tan, Manxiu Ma, Yu Fu, Donghui Wu, Guozhi Luo, Ping Ren

**Author notes:** **Correspondence to:** Ping Ren; Address: No. 1080 Cuizhu Rd., Shenzhen, Guangdong, China.

## Abstract

Risky decision-making under uncertainty reflects complex cognitive processes supported by distributed brain networks that are vulnerable to aging. However, it remains unclear whether risk-taking behavior can serve as a behavioral marker of brain aging. In the present study, we combined behavioral tasks, computational modeling, and structural magnetic resonance imaging to investigate the relationship between risky decision-making, chronological age, and brain age. A total of 55 young adults and 112 healthy older adults completed the Iowa Gambling Task (IGT) and the Balloon Analogue Risk Task (BART), along with neuropsychological assessments and neuroimaging scanning. Decision processes were quantified using computational models, including the Value-Plus-Perseveration model and Exponential-Weight Mean–Variance. Brain age was estimated from gray matter volume. The results showed significant age-related alterations in parameters reflecting feedback sensitivity, learning rate, and loss aversion in both tasks. Within older adults, several decision parameters were significantly associated with both chronological age and brain age. Regression analyses further showed that computational parameters significantly predicted chronological age and brain age, whereas traditional cognitive screening measures did not show significant predictive effect. Structural brain analyses indicated that IGT-related parameters were primarily associated with the basal ganglia, while BART-related parameters were linked to a broader network including prefrontal, cingulate, and temporal regions. These findings suggest that computational markers of risk-taking behavior capture subtle age-related changes in cognitive processes and brain deterioration. Therefore, risk-taking parameters may serve as reliable functional markers of brain aging, providing critical insights into the mechanisms underlying successful aging.

## Introduction

Decision-making under risk is a critical ability that enables individuals to adapt to complex and uncertain environments. This ability relies on cognitive function in multiple domains, including valuation, learning, memory, and affective processes, supported by distributed cortical and subcortical brain networks [1]. Accumulating evidence indicates that risky decision-making is particularly vulnerable to aging, even among older adults with normal cognitive scores in neuropsychological assessments [2, 3]. Importantly, age-related alterations in risky decision-making often emerge before overt cognitive impairment [4], suggesting that such behaviors may capture subtle changes in brain health that are not detectable by conventional cognitive screening tools.

Despite growing interest, findings on age-related differences in risky decision-making remain mixed [5, 6]. The Iowa gambling task (IGT) and balloon analogue risk task (BART) are among the most widely used paradigms to assess decision-making under uncertainty. The IGT captures learning-based decision-making under uncertainty implicitly, requiring individuals to maximize their monetary rewards through distinguishing advantageous and disadvantageous options [7, 8]. In contrast, the BART measures a more direct, moment-to-moment propensity to take explicit risk for reward [9, 10]. Although both the two tasks are popular paradigms for assessing risk-taking behavior, they engage partially distinct cognitive and motivational components. Previous studies comparing the IGT and BART have shown mixed findings, with some reporting significant relationships [11–13], while others found no relationship at all [14–16]. These inconsistencies suggest that traditional behavioral measures may be insufficiently sensitive to underlying changes in decision processes. Moreover, age-related alterations in risky decision-making have been found more complicated and conflicting as well, highlighting substantial inter-individual variability in aging trajectories [17]. Therefore, latent decision mechanisms across tasks would be required to clarify its relationship with chronological aging and cognitive integrity.

Recent advances in computational modeling provide a powerful framework for decomposing complex decision behavior into interpretable cognitive parameters [18]. For the IGT, the Value-Plus-Perseveration (VPP) model disentangles value-based learning from choice perseveration, offering improved explanatory power over traditional reinforcement learning models [19]. A growing number of studies have shown that the VPP model successfully characterizes altered decision-making processes in various clinical or neuropsychological conditions [20–22]. For the BART, the Exponential-Weight Mean-Variance (EWMV) model integrates dynamic belief updating and risk sensitivity, overcoming limitations of earlier computational models [23]. These computational approaches have been successfully applied to characterize altered decision-making in psychiatric and neurological conditions [20–22, 24]. However, it is still unclear whether computational parameters of risky decision-making can serve as sensitive markers of age-related cognitive and brain changes.

In parallel, the concept of “brain age”, typically estimated from structural magnetic resonance imaging (MRI) features such as gray matter volume, has been found to be a promising neuroimaging-based biomarker of brain health [25]. For example, a large brain-age gap (discrepancy between estimated brain age and chronological age) has been associated with poorer cognitive performance and increased risk for neurodegenerative conditions [26]. In contrast, widely used neuropsychological screening tools, such as the Mini-Mental State Examination (MMSE) and Montreal Cognitive Assessment (MoCA), show limited sensitivity to subtle or preclinical cognitive decline and weak associations with brain structural measures in healthy aging populations [27, 28]. Notably, risky decision-making has been supported by large-scale neural networks involving the prefrontal cortex, anterior cingulate cortex, insula, and striatum, which are particularly susceptible to age-related structural and functional decline [29]. So far was we know, there has been less study on the relationships between risk-taking behavior and chronological age or brain age. We speculate that risky decision-making tasks, especially when quantified using computational models, may offer a unique window into brain aging by linking cognitive processes to their neural substrates.

To address these questions, the present study integrated behavioral measures, computational modeling, and structural MRI data to characterize the cognitive and neural mechanisms of altered risky decision-making processes in aging. By comparing conventional neuropsychological tools and risky decision-making tasks, we further investigated their effects in estimating chronological and brain age in older adults. The aims of this study were three-folds: 1) Whether computational modeling parameters would provide greater sensitivity to aging-related alterations in risky decision-making than conventional behavioral indices. 2) To investigate the task-specific decision mechanisms (IGT vs. BART) are differentially affected by aging. 3) Whether risk-taking behavior would be more informative in estimating chronological age and brain age relative to conventional cognitive screening assessments (i.e., MOCA, MMSE).

## Material and Methods

### Participants

Totally, sixty-eight young adults (aged 19 – 31 years) and 129 older adults (aged 54 – 88) were recruited from multiple communities in Shenzhen (**Figure 1**). The individuals with one of the following conditions were excluded in the following analysis: (1) a history of diagnosed neurological or psychiatric diseases (i.e., major depression, anxiety, or cerebrovascular disease); (2) MRI contraindications (i.e., metallic implant, claustrophobia, or pacemaker). (3) participants with left handedness were excluded to avoid lateralization of brain deterioration. Each participant was required to sign a written informed consent form after a full written and verbal explanation of the study. This study was performed in accordance with the Declaration of Helsinki, and had been approved by the Ethics Committee of Shenzhen Kangning Hospital.

**Figure 1.**
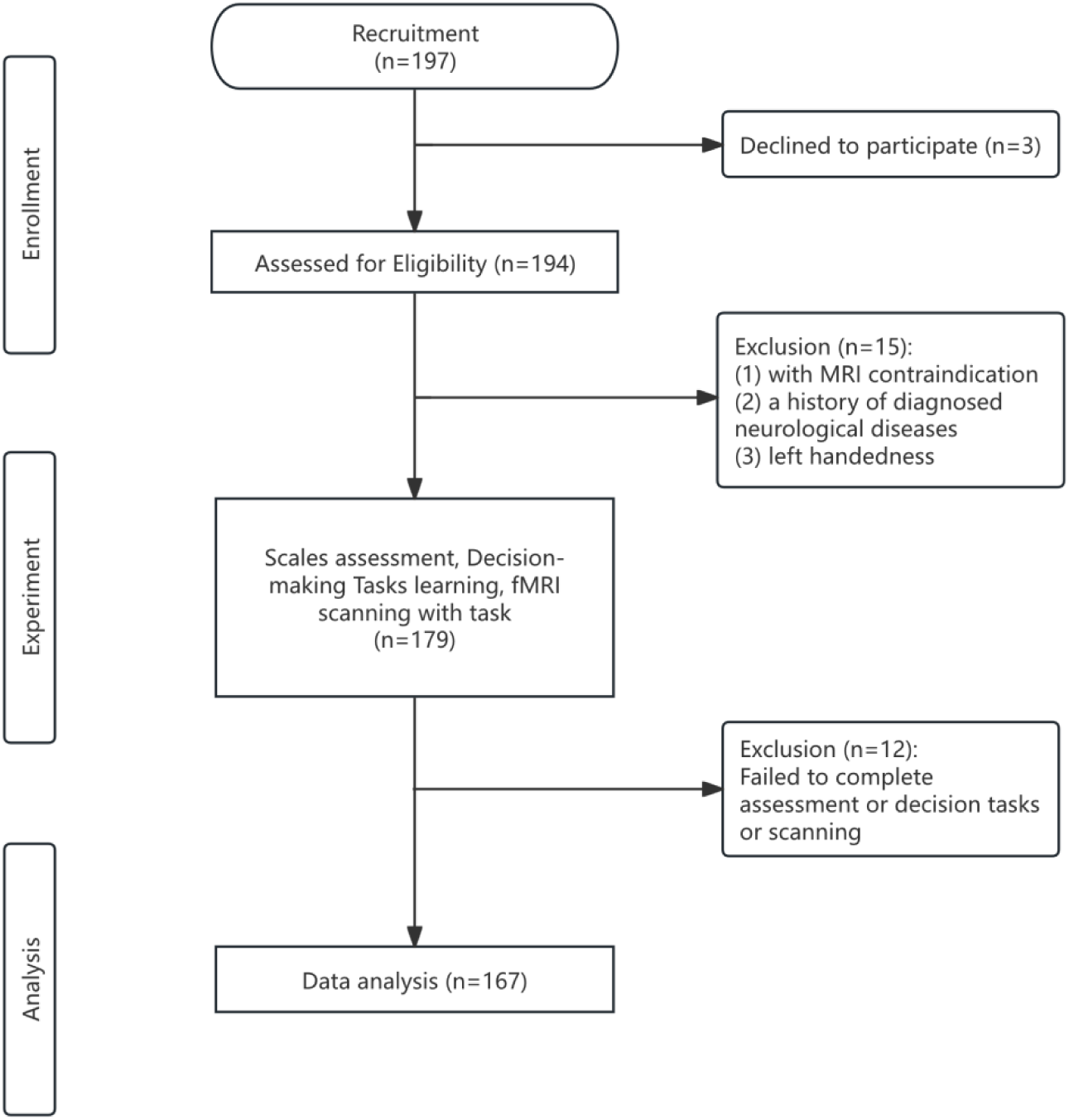
**Flow chart of participant enrolment.**

For each participant, global cognitive integrity was assessed using the MoCA and MMSE. The MMSE is a 30-point questionnaire that is widely used in clinical and research settings to assess cognitive function in multiple domains, including orientation, short-term memory, spatial abilities and so on [30]. The MoCA is a popular screening tool for assessing global cognitive ability, showing high sensitivity in detecting cognitive impairment [31, 32]. To better examine the relationships between cognitive state and other variables, the current analyses included participants with a broad range of MoCA (20 – 30) and MMSE (16 – 29).

### Risky decision-making tasks

The modified IGT was applied based on Cauffman and colleague’s version [33], which has been successfully used in our previous studies [34, 35]. In each trial, participants were required to ‘play’ or ‘pass’ the target card highlighted with a yellow frame randomly. If choosing ‘play’, they would win or lose monetary reward (a net gain/loss) with the intrinsic possibility. Therefore, choosing from the advantageous decks (deck C and D) would give participants a cumulative gain, while choosing from the disadvantageous decks (deck A and B) would give a loss. With a banked amount of ¥100 at the beginning of the IGT, participants were required to earn monetary reward as much as possible. For each participant, a net score (advantageous versus disadvantageous selections) was computed to measure risk-taking behavior in data analysis.

In the BART, participants were presented with a colored balloon (red or blue) and required to press the button in their right hand to pump the balloon, and press the button in their left hand to collect monetary earnings. With the balloon inflating, monetary earnings in the “wallet” were increased, as well as the risk of explosion probability. Money in the “wallet” would be successfully transferred to the “bank” if participants press stop pumping before explosion, otherwise the money in the “wallet” would be lost. Two colored balloons indicated different explosion probability: red balloon with a high risk of explosion and blue balloon with a low risk of explosion. The proportion of successful trials and adjusted number of pumps were used to measure risk-taking behavior the same as in the previous studies [9, 36].

### Computational modeling analyses

To comprehensively quantify decision-making processes during task performance, our behavioral data were fitted to computational models of the IGT and BART, separately (**Table 2**). For the IGT, we tested the VPP model and other three popular models: the Outcome-Representation Learning model (ORL) [37], the Prospect Valence Learning model with the Delta rule (PVL-Delta) [38], the Prospect Valence Learning model with Decay (PVL-Decay) [39]. For the BART, we compared the EWMV model and 4-parameter model [40]. Detailed descriptions of each model are provided in the **Supplementary Materials**. Model comparisons were conducted using the Leave-One-Out Information Criterion (LOOIC) and the Widely Applicable Information Criterion (WAIC). For each task, the best fitted model was used in the following analysis.

### Imaging data acquisition and analyses

Structural MRI data were acquired on a 3.0T MRI system (Discovery MR750 System, GE Healthcare) with an eight-channel phased-array head coil. The T1-weighted structural images were acquired by using a three-dimensional brain volume imaging sequence that covered the whole brain (repetition time (TR) = 6.7 ms, echo time (TE) = 2.9 ms, flip angle = 7 degrees, matrix = 256×256, slice thickness = 1 mm, 196 slices). The images were preprocessed in the DPABI_v4.3 (https://rfmri.org/dpabi) based on the SPM12 (http://www.fil.ion.ucl.ac.uk/spm/) [41]. Then participants were required to complete the IGT and BART during a task-related fMRI session. In the present study, we only used task behavioral data, and the task-related imaging data were analyzed and reported elsewhere.

### Gray matter atrophy and brain age

The structural images were preprocessed to generate a whole-brain gray matter map using Voxel-based morphometry (VBM). For each participant, the T1-weighted image was segmented into gray matter, white matter, and cerebrospinal fluid. Then, a gray matter template was generated through an iterative nonlinear registration using DARTEL, a toolbox with a fast diffeomorphic registration algorithm [42]. For each participant, an averaged gray matter (GM) volume of the entire brain was computed to control age-induced brain atrophy in the following analyses. Then the GM images were resampled to 3 × 3 × 3 mm, and smoothed with a Gaussian kernel (FWHM = 6 mm). Based on the Automated Anatomical Labeling atlas (AAL) [43], averaged GM values within 90 brain regions were selected to calculate individual’s brain age. A generalized linear model was used to predict brain age by using GM values as variables.

### Statistical analysis

Other statistical analyses were conducted in the Matlab 2019b and SPSS V22. The independent t-test/Chi-square (*χ^2^*) test was used to examine the group differences in demographic, neuropsychological measurements and task performance between young and older adults. The nonparametric test (Mann-Whitney U test) was used to examine the group differences in behavioral parameters both in IGT and BART. Then a partial correlation was used to examine the relationships between behavioral parameters and chronological age/brain age. Linear regression models were used to assess the relationship between chronological/brain age (outcome) and neuropsychological measurements (predictors), behavioral parameters (predictors), adjusting for gender and education. In the gray matter analysis, the Pearson correlation was used to examine the association between Behavioral parameters and GMV derived from 90 regions in the AAL. To comprehensively exhibit all possible significant correlations, uncorrected p < 0.05 was used as a statistical threshold without correcting multiple comparisons.

## Results

### Demographic, cognitive measurements and task behavioral analyses

Eventually, 55 young adults (aged 19 – 31 years) and 112 older adults (aged 54 – 88) were included in the following analyses (**Figure 1**). Compared with young adults, older adults exhibited significantly lower level of education, reduced MMSE and MoCA scores. No behavioral difference was observed in the IGT or BART (**Table 1**).

**Table 1.** Demographic data, cognitive assessment and task performance.

| | Mean ± SD | | $t / \chi^2$ | **p** |
| --- | --- | --- | --- | --- |
|  | Young (n = 55) | Older (n = 117) |  |  |
| age | 24.55 ± 2.90 | 64.02 ± 5.90 | -58.84 | < 0.001 |
| gender | 29/26 | 50/67 | 1.50 | 0.220 |
| education | 13.52 ± 2.75 | 10.32 ± 2.62 | 7.35 | < 0.001 |
| MMSE | 28.98 ± 1.58 | 27.57 ± 2.31 | 4.67 | < 0.001 |
| MoCA | 26.95 ± 1.96 | 22.03 ± 3.48 | 11.80 | < 0.001 |
| *IGT* |  |  |  |  |
| Good selection (%) | 0.81 ± 0.13 | 0.80 ± 0.11 | 0.02 | 0.892 |
| Bad selection (%) | 0.73 ± 0.13 | 0.75 ± 0.14 | 0.46 | 0.498 |
| Net score | 0.08 ± 0.19 | 0.05 ± 0.15 | 0.23 | 0.630 |
| *BART (high-risk)* |  |  |  |  |
| successful trials (%) | 0.69 ± 0.12 | 0.73 ± 0.15 | 0.02 | 0.887 |
| adjusted pumps | 2.60 ± 0.45 | 2.38 ± 0.63 | 0.04 | 0.835 |
| *BART (low-risk)* |  |  |  |  |
| successful trials (%) | 0.79 ± 0.11 | 0.88 ± 0.11 | 2.16 | 0.143 |
| adjusted pumps | 4.39 ± 0.66 | 3.50 ± 1.16 | 2.10 | 0.149 |

Note. The independent t-test/Chi-square ( $\chi^2$ ) test was used to examine the group differences in demographic and neurocognitive measurements. The non-parametric test (Mann-Whitney U test) was used to examine the group differences in behavioral indices in the IGT and BART. In the BART, high- and low-risk refer to the red- and blue-balloon conditions, respectively. Abbreviations, MoCA, Montreal Cognitive Assessment; MMSE, Mini-Mental State Examination.

### Modeling analysis in the IGT and BART

In addition to task behavioral metrics, modeling analyses were used to comprehensively characterize risk-taking behavior in the young and older groups. Based on the model comparisons, the VPP model provided the best fitness for the data in IGT, while the EWMV model demonstrated superior performance in BART (**Table 2**). Therefore, the VPP and EWMV models were selected to do the following analyses. **Table 3** exhibited modeling parameters of the two tasks in young and older adults. In the IGT, older adults exhibited significant changes in multiple parameters: higher values in the A (memory decay), lambda (loss aversion) and K (perseverance); lower values in alpha (feedback sensitivity), cons (consistency), epP (gains impact), epN (losses impact), and w (reinforcement-learning rate). In the BART, the high- (red balloon) and low-risk (blue balloon) conditions were computed separately. In the high-risk condition, older adults showed significantly increased value in lambda (loss aversion), while decreased values in phi (prior belief of burst) and eta (learning rate). In the low-risk condition, older adults exhibited significantly increased lambda and decreased eta. Within each model, FDR corrected p < 0.05 was used for multiple comparisons.

**Table 2.** Computational models in the IGT and BART.

|  | **Young** |  | **Older** |  |
| --- | --- | --- | --- | --- |
|  | LOOIC | WAIC | LOOIC | WAIC |
| *IGT model* |  |  |  |  |
| VPP | 9494.992 * | 9492.998 * | 19461.90 * | 19450.95 * |
| ORL | 9560.580 | 9558.007 | 19626.68 | 19611.08 |
| PVL_decay | 9559.938 | 9567.852 | 19682.81 | 19680.79 |
| PVL_delta | 9557.178 | 9561.412 | 19695.85 | 19694.79 |
| *BART model* |  |  |  |  |
| EWMV | 2713.240 * | 2626.850 * | 4773.573 * | 4627.790 * |
| 4-Parameters model | 2813.679 | 2722.308 | 4808.863 | 4656.731 |

Note. lower LOOIC and WAIC values indicate a better fit to the data. Abbreviations, EWMV, Exponentially-Weighted Mean-Value; LOOIC, Leave-One-Out Information Criterion; ORL, Outcome-Representation Learning; PVL, Prospect Valence Learning; WAIC, Applicable Information Criterion; VPP, Value-Plus-Perseverance. \*, best performance among the models.

**Table 3.** Model parameters in the IGT and BART.

|  | Median |  | z | p |
| --- | --- | --- | --- | --- |
|  | Young | Older |  |  |
| *VPP (IGT)* |  |  |  |  |
| A | 0.03 | 0.08 | -9.77 | < 0.001 |
| alpha | 1.10 | 0.33 | -10.49 | < 0.001 |
| cons | 2.21 | 0.72 | -10.49 | < 0.001 |
| lambda | 3.65 | 3.80 | -10.33 | < 0.001 |
| epP | -1.19 | -1.35 | -5.01 | < 0.001 |
| epN | -0.62 | -0.75 | -10.48 | < 0.001 |
| K | 0.91 | 0.94 | -9.39 | < 0.001 |
| w | 0.95 | 0.79 | -10.49 | < 0.001 |
| *EWMV (BART high-risk)* |  |  |  |  |
| phi | 0.07 | 0.01 | -10.48 | < 0.001 |
| eta | 0.03 | 0.00 | -10.34 | < 0.001 |
| rho | 0.00 | -0.00 | -0.57 | 0.567 |
| tau | 9.32 | 8.11 | -1.67 | 0.096 |
| lambda | 4.68 | 33.22 | -10.49 | < 0.001 |
| *EWMV (BART low-risk)* |  |  |  |  |
| phi | 0.03 | 0.03 | -1.35 | 0.176 |
| eta | 0.01 | 0.01 | -5.19 | < 0.001 |
| rho | -0.30 | -0.02 | -0.86 | 0.201 |
| tau | 7.16 | 8.20 | -1.10 | 0.270 |
| lambda | 1.98 | 10.36 | -10.28 | < 0.001 |

**Note.** The Mann-Whitney U test was applied to compare model parameters between the young and older groups (FDR corrected $p < 0.05$ ). The *z*-value reflects group difference in rank distributions with larger absolute value indicating greater difference. Abbreviations, EWMV, Exponentially-Weighted Mean-Value; VPP, Value-Plus-Perseverance.

### Risk-taking behavior and chronological age in older adults

Within the older group, Spearman correlation analysis was applied to examine the relationships between chronological age and model parameters in the IGT and BART. In the IGT, chronological age was negatively correlated with alpha and k, while positively correlated with epP and w. In the BART high-risk condition, chronological age was positively correlated with eta and negatively correlated with tau (**Figure 2**). No significant correlation was observed in the BART low-risk condition (**Supplementary Table 1**).

**Figure 2.**
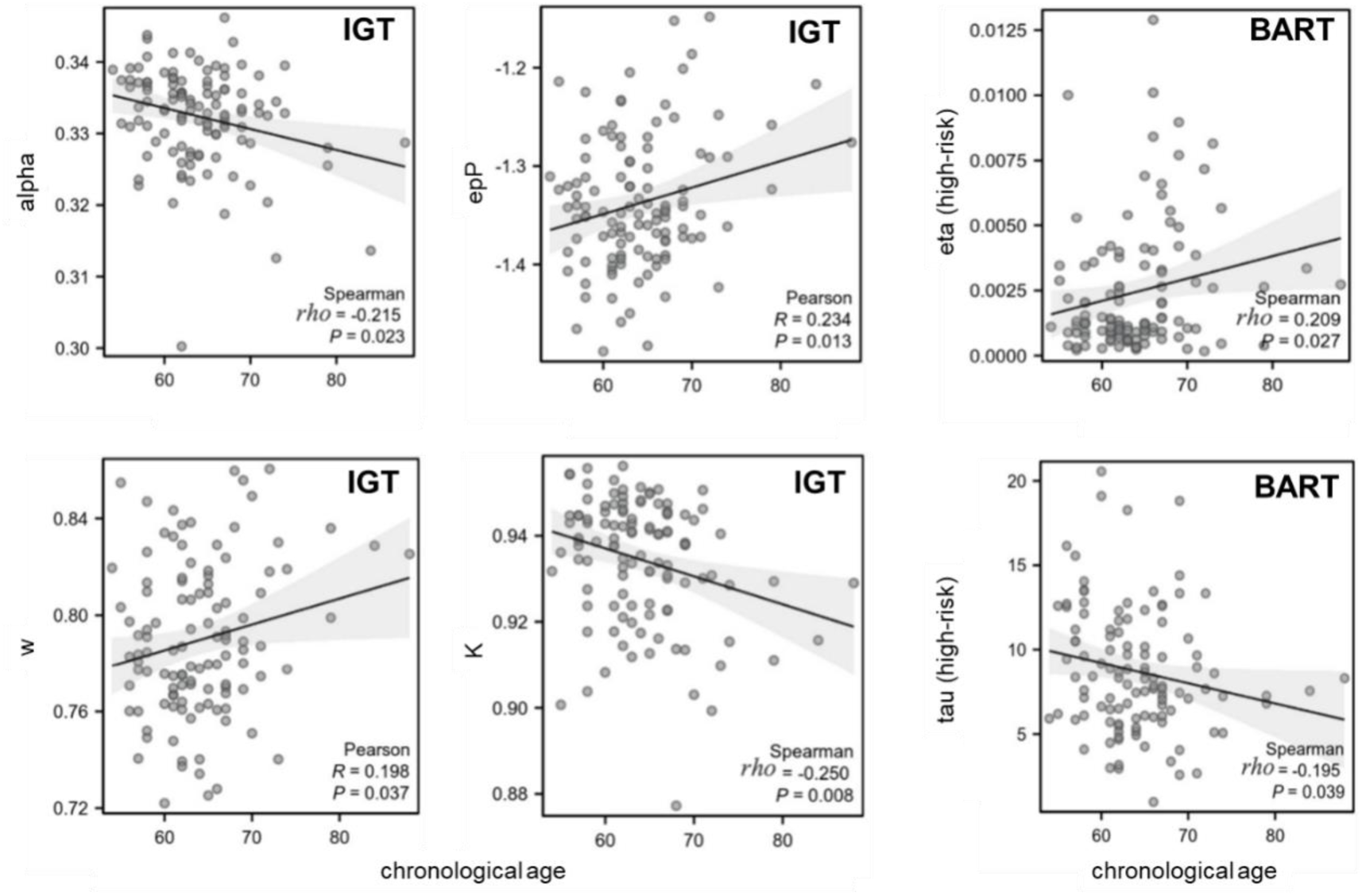
Correlations between model parameters and chronological age in older adults. In the IGT, chronological age was found negatively correlated with alpha and K, while positively correlated with epP and w. In the BART, chronological age was significantly correlated with parameters under the high-risk condition, showing negative correlation with eta and positive correlation with tau. All regression models were controlled for gender and years of education.

According to the correlation analysis, the parameters significantly correlated to chronological age were selected as predictors in the linear regression model for the IGT and BART (**Table 4, Supplementary Figure 1**).

**Table 4.** Linear regression models in predicting chronological age in older adults.

| Input | B | SE | $\beta$ | t | p |
| --- | --- | --- | --- | --- | --- |
| *Model 1* |  |  |  |  |  |
| MMSE | -0.347 | 0.288 | -0.138 | -1.208 | 0.230 |
| MoCA | -0.159 | 0.199 | -0.094 | -0.798 | 0.427 |
| *Model 2 (IGT)* |  |  |  |  |  |
| alpha | -241.690 | 103.777 | -0.279 | -2.329 | 0.022 |
| epP | -16.395 | 25.761 | -0.188 | -0.636 | 0.526 |
| K | -122.330 | 67.744 | -0.302 | -1.806 | 0.074 |
| w | 10.739 | 49.165 | 0.058 | 0.218 | 0.828 |
| *Model 3 (BART high-risk)* |  |  |  |  |  |
| eta<sub>high-risk</sub> | 428.121 | 221.498 | 0.181 | 1.933 | 0.056 |
| tau<sub>high-risk</sub> | -0.274 | 0.148 | -0.173 | -1.850 | 0.067 |
| *Model 4 (IGT + BART)* |  |  |  |  |  |
| alpha | -256.227 | 102.237 | -0.296 | -2.506 | 0.014 |
| epP | -25.842 | 25.444 | -0.297 | -1.016 | 0.312 |
| K | -118.476 | 66.239 | -0.293 | -1.789 | 0.077 |
| w | 22.897 | 48.324 | 0.124 | 0.474 | 0.637 |
| eta<sub>high-risk</sub> | 441.800 | 215.695 | 0.187 | 2.048 | 0.043 |
| tau<sub>high-risk</sub> | -0.183 | 0.146 | -0.116 | -1.255 | 0.212 |

**Note.** The model parameters, showing significant correlations with chronological age, were involved as predictors in Model 2, 3, and 4. All regression models were controlled for gender and years of education. Abbreviations, MoCA, Montreal Cognitive Assessment; MMSE, Mini-Mental State Examination.

Model 1: chronological age = β^1^MMSE + β^2^MOCA + covariates + ε.
Model 2 (IGT): chronological age = β^1^alpha + β^2^epP + β^3^K + β^4^w + covariates + ε.
Model 3 (BART high-risk): chronological age = β^1^eta^high-risk^ + β^2^tau^high-risk^ + covariates + ε.
Model 4 (IGT + BART): chronological age = β^1^alpha + β^2^epP + β^3^K + β^4^w + β^5^eta^high-risk^ β^6^tau^high-risk^ + covariates + ε.

In specific, Model 1 showed no significant predictive power using MOCA and MMSE as predictors (F=2.140, p=0.123, R^2^=0.037). Model 2 (predictors: IGT model parameters) showed significant predictive power (F=3.621, p=0.008, R^2^=0.117). Model 3 (predictors: BART high-risk parameters) also showed significant predictive power (F=4.434, p=0.014, R^2^=0.074). Notably, Model 4 (combining IGT and BART parameters) exhibited the strongest predictive power (F=3.269, p=0.002, R^2^=0.171), in which feedback sensitivity (alpha, β=-0.296, t=-2.506, p=0.014) and updating exponent (eta ^high-risk^, β=0.187, t=2.048, p=0.043) showed significant effect on chronological age.

### Risky decision-making and brain age in older adults

In older adults, estimated brain-age was calculated based on gray matter volumes derived from 90 AAL brain regions using a linear regression model. In the IGT, correlation analyses showed that estimated brain age was significantly associated with alpha, epP, and k (**Figure 3A**). In the BART high-risk condition, eta was positively correlated with brain age (**Figure 3B**). No significant correlation was observed in the BART low-risk condition (**Supplementary Table 2**).

**Figure 3.**
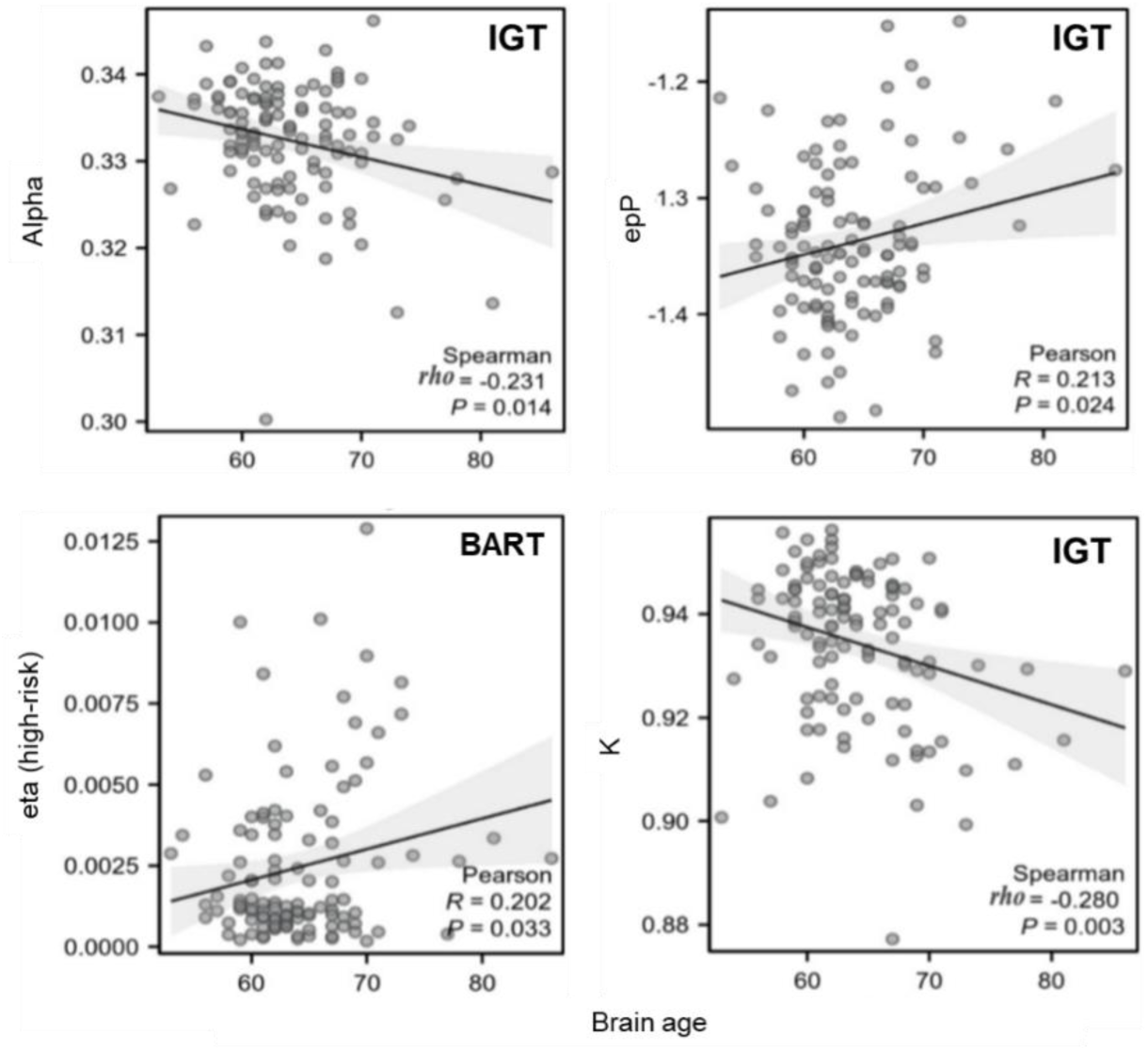
Correlations between model parameters and brain age in older adults. In the IGT, Brain age was found negatively correlated with alpha and K, while positively correlated with epP. In the BART, brain age showed significant correlation with eta under the high-risk condition. All regression models were controlled for gender and years of education.

According to the correlation analysis, the parameters significantly correlated to brain age were selected as predictors in the linear regression model for the IGT and BART (**Table 5, Supplementary Figure 2**).

**Table 5.**
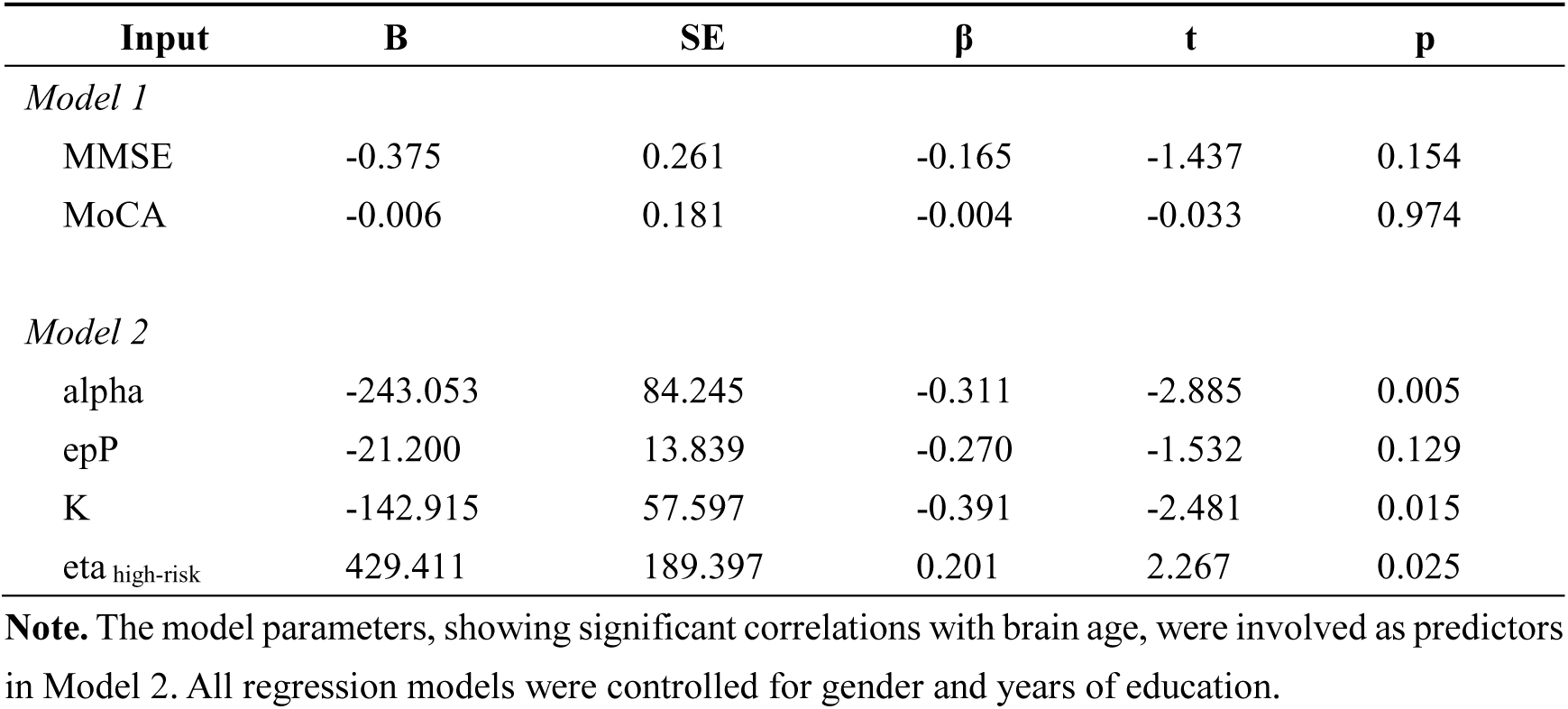
Linear regression models in predicting brain age in older adults.

Model 1: brain age = β^1^MMSE + β^2^MOCA + covariates + ε.
Model 2 (IGT+BART): brain age = β^1^alpha + β^2^epP + β^3^K + β^4^eta^high-risk^ + covariates + ε.

In specific, Model 1 had no significant predictive power using MOCA and MMSE as predictors (F=1.459, p=0.237, R^2^=0.026). Combining the IGT and BART parameters, Model 2 showed significantly predictive power (F=5.347, p=0.001, R^2^=0.163). Specifically, Feedback sensitivity (alpha, β=-0.311, t=-2.885, p=0.005), Perseverance decay (K, β=-0.391, t=-2.481, p=0.015) and Updating exponent (eta ^high-risk^, β=201, t=2.267, p=0.025).

### Regional brain atrophy and risky decision

In the GM analysis, GM volumes derived from 90 brain regions were correlated to model parameters, behavioral measures and neuropsychological assessments. There was no significant correlation observed between GM and MMSE/MOCA (**Figure 4A-D, Supplementary Figure 3A**). In the IGT, risk aversion (lambda) showed negatively significant correlation with GM in the caudate, putamen, pallidum, and superior occipital gyrus (**Figure 4E-H, Supplementary Figure 3B**). In the BART high-risk condition, prior belief of burst (phi) showed positive correlations with GM, while risk preference (rho) showed negative correlations with GM in multiple brain regions (**Figure 5A-D, Supplementary Figure 4A**). In the BART low-risk condition, prior belief of burst (phi) and Risk preference (rho) showed negative correlation, while inverse temperature (tau) and loss aversion (lambda) showed positive correlation in multiple brain regions (**Figure 5E-H, Supplementary Figure 4B**).

**Fig 4.**
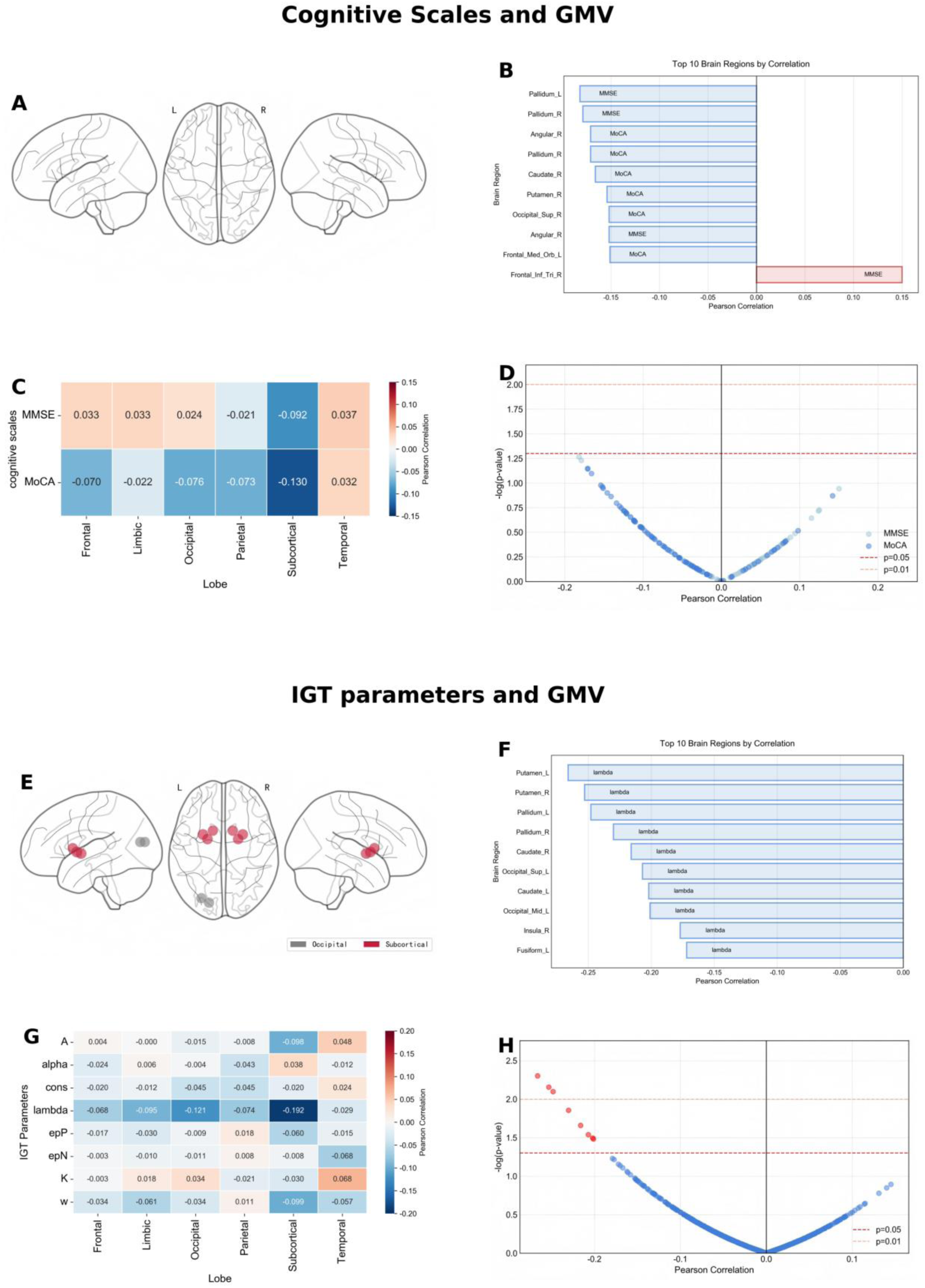
Correlations between cognitive scores/IGT and GM. A-D show MMSE and MoCA correlations: (A) brain regions with significant correlations (p < 0.05), (B) top 10 brain regions in correlation analysis, (C) average correlations across brain areas, (D) volcano plot displaying statistical significance versus effect size (Pearson correlation) with threshold lines at p = 0.05 and p = 0.01. E-H illustrate IGT parameter-GM correlations.

**Fig 5.**
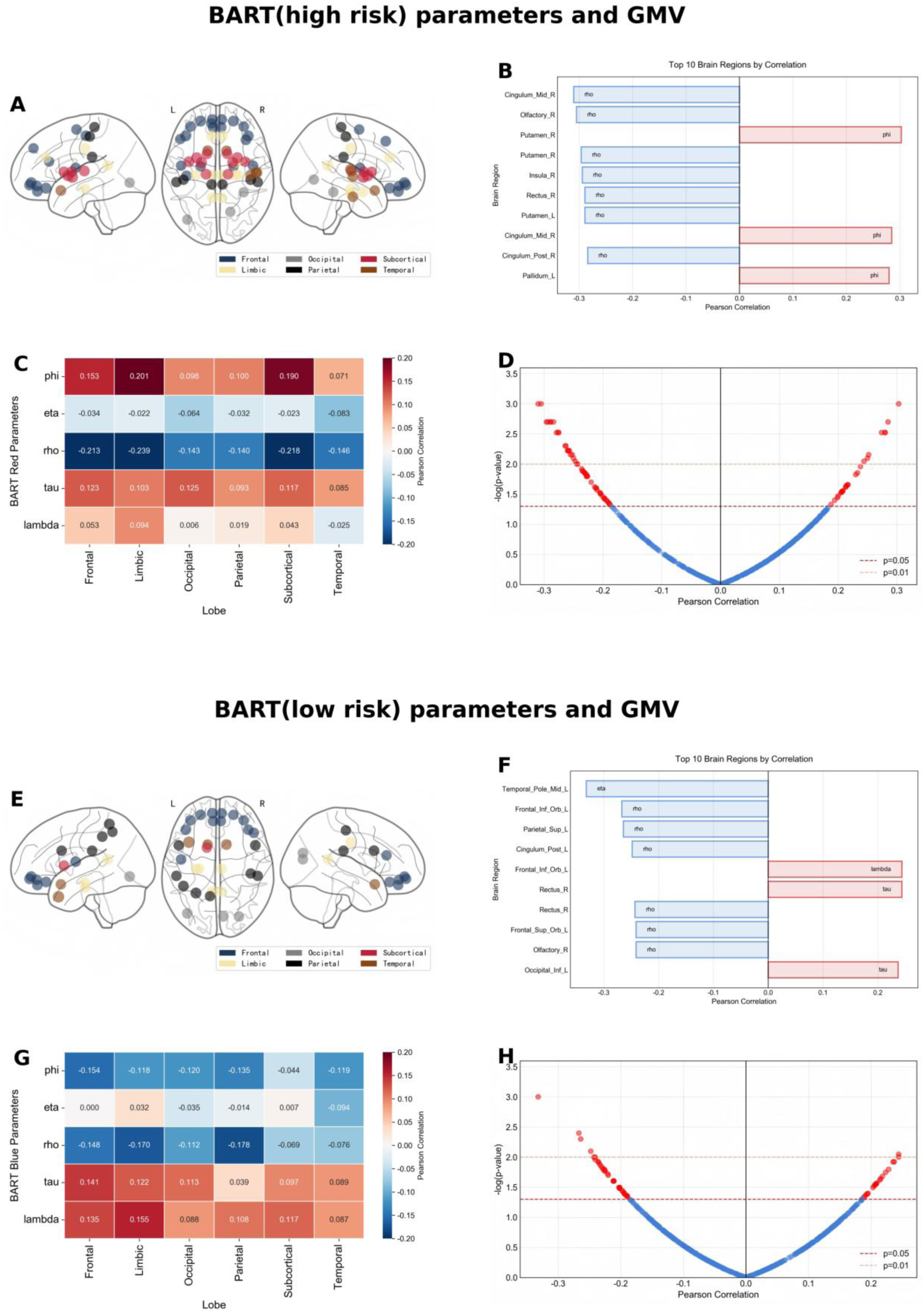
Correlations between BART parameters and GM. A-D show BART high-risk condition: (A) brain regions with significant correlations (p < 0.05), (B) top 10 brain regions in correlation analysis, (C) average correlations across brain regions, (D) volcano plot displaying statistical significance versus effect size (Pearson correlation) with threshold lines at p = 0.05 and p = 0.01. E-H illustrate BART low-risk condition.

## Discussion

The present study investigated the relationship between risky decision-making, chronological age, and brain age by integrating behavioral paradigms, computational modeling, and structural MRI. Although older adults showed no significant difference from young adults in conventional behavioral indices from risky decision-making tasks, computational modeling revealed clear age-related alterations in multiple decision parameters. Importantly, several of these parameters were significantly associated with both chronological age and MRI-derived brain age in older adults. Regression analyses further demonstrated that risk-taking parameters significantly predicted chronological age and brain age, whereas traditional cognitive screening measures showed no significant predictive power. In addition, brain–behavior analyses indicated that IGT-related parameters were mainly associated with basal ganglia, whereas BART-related parameters were linked to a broader network. So far as we know, this is the first study that systematically investigated the differences of cognitive and neural mechanisms between the IGT and BART, and highlight the potential of computational markers of risky decision-making in predicting brain aging.

In conventional behavioral analysis, older adults did not differ from young adults in the IGT and BART, in line with previous studies [44, 45]. However, computational modeling analyses revealed age-related differences by decomposing decision-making processes into latent components, including learning, memory, feedback sensitivity, and choice persistence. In the IGT, older adults exhibited a higher memory decay parameter (A), indicating faster forgetting of past outcomes and reduced integration of accumulated experiences. This pattern is consistent with well-established age-related declines in working memory, which may limit the ability to maintain and integrate value information across trials. Older adults also showed greater loss aversion (lambda), reflecting a stronger tendency to avoid potential losses and a more risk-averse decision profile in later life. In addition, older adults demonstrated reduced feedback sensitivity (alpha) and a lower reinforcement-learning rate (w), suggesting less efficient updating of value representations based on experienced outcomes. Together, these changes indicate that aging may shift IGT performance away from stable, magnitude-sensitive learning toward strategies that are less responsive to outcome values and more constrained by memory and updating limitations. Moreover, older adults showed increased perseveration (K) and reduced gain and loss sensitivity (epP/epN), further suggesting greater cognitive rigidity and diminished value sensitivity in aging. For the BART, the EWMV model revealed a similar motivational shift. Older adults exhibited greater loss aversion (λ) in both high- and low-risk conditions, suggesting that potential negative outcomes exert a stronger suppressive effect on risk-taking behavior. This finding is consistent with theories proposing increased harm avoidance or greater sensitivity to negative feedback in later life [46]. Moreover, older adults exhibited reduced belief updating (eta), implying a lower rate of integrating new outcome information into beliefs about the probability of balloon explosion. Lower eta is compatible with reduced cognitive flexibility and could manifest behaviorally as slower adaptation to changing risk levels or reduced use of recent information to calibrate pumping behavior. In the high-risk condition, older adults also showed lower phi (prior belief of burst), indicating different initial expectations about risk. Taken together, these parameters point to a decision profile in aging characterized by altered value sensitivity, reduced learning efficiency and cognitive flexibility.

Notably, conventional neuropsychological screening tools such as the MMSE and MoCA have been shown to be relatively insensitive to subtle, preclinical cognitive alterations associated with aging [47, 48]. Consistent with previous findings, our results indicated that neither MMSE nor MoCA exhibited significant predictive power for chronological age or brain age in the regression analyses. In contrast, computational parameters derived from the IGT and BART demonstrated significant predictive value for both chronological age and MRI-derived brain age. These results support our hypothesis that risk-taking behavior under uncertainty, particularly when quantified using computational modeling, may capture subtle and multidimensional cognitive changes that are not detectable using global cognitive screening measures. Importantly, individuals whose estimated brain age exceeds their chronological age are often considered to exhibit accelerated brain aging, which has been associated with increased risk for cognitive decline and adverse health outcomes [49, 50]. Our findings therefore suggest that decision-making parameters may serve as accessible behavioral correlates of brain aging and may provide a complementary approach for monitoring brain health in older adults. Moreover, because computational parameters reflect specific cognitive processes, they may help identify which components of decision-making (e.g., feedback sensitivity, loss aversion, or learning efficiency) are most closely linked to brain aging, thereby providing potential targets for future interventions.

Crucially, the current study extends these behavioral observations to anatomical changes of aging brain. ROI-based gray matter analyses indicated that IGT- and BART-derived parameters are associated with distinct brain regions, confirming that the two tasks rely on different neural mechanisms. In specific, IGT parameters were predominantly associated with GM in the striatum (i.e., caudate, putamen, pallidum), consistent with the role of the striatum in habit formation and implicit reinforcement learning [51, 52]. While the BART engaged a more widespread distributed network involving the prefrontal cortex, insula, and cingulate gyrus, which requires explicit, trial-by-trial risk calculation and inhibitory control [24]. Therefore, the distinct correlations between task-specific parameters and gray matter imply that computational components are not merely abstract concepts of behavior but reflect underlying neurobiological variation. Interestingly, we didn’t observe significant correlations between GM and MMSE/MoCA in the older group. This finding is consistent with previous neuroimaging studies showing that global cognitive screening measures may remain stable despite structural brain changes in healthy aging populations, reflecting the limited sensitivity and ceiling effects of these instruments in detecting subtle neural alterations [53, 54]. Compared with conventional neuropsychological screening tools, decision-making parameters derived from computational modeling may provide more sensitive markers of brain health in aging. By decomposing risk-taking behavior into interpretable components such as learning efficiency, feedback sensitivity, and loss aversion, computational parameters offer mechanistic insights into specific cognitive–affective processes that may be particularly vulnerable to age-related brain alterations. Integrating such computational markers with traditional cognitive assessments may therefore improve the characterization and monitoring of brain aging.

Several limitations need to be acknowledged. First, the study design is cross-sectional, therefore, we cannot determine the causality between risk-taking alterations and brain aging. Longitudinal studies will be necessary to test whether baseline decision parameters predict future brain-age acceleration or cognitive decline. Second, the current study only involved the IGT and BART to examine the relationships between risk-taking behaviors and chronological age/brain age. It should be cautious to generalize the conclusions to other risky decision-making tasks. Third, there have been several methods to predict brain age in previous studies, showing distinct accuracy and test-retest liability [55]. Here, although we found consistent results by using generalized linear model, it is worthy to retest the relationship by using other brain age prediction models.

## Conclusion

In summary, this study provides evidence that computational modeling parameters of risky decision-making capture aging-related changes that are not apparent in traditional behavioral metrics, and that these features relate to both chronological age and MRI-derived brain age. The findings highlight that risk-taking parameters are not only meaningful behavioral markers, but also robust indicators of brain health in older adults, providing novel insights on the early detection and monitoring of neurocognitive aging.

## Acknowledgements

We appreciate all who participated in this study. This work was supported by: Natural Science Foundation of Guangdong Province 2023A1515012911 (to PR), Shenzhen Science and Technology Innovation Commission JCYJ20210324133208023 (to PR), Sanming Project of Medicine in Shenzhen SZSM202311026, National Natural Science Foundation of China 82360271 (to YF), and Shenzhen Fund for Guangdong Provincial High-level Clinical Key Specialties SZGSP013.

## Data Availability Statement

The data used in this study is available from the corresponding author upon request.

## References

1. Meshulam, L., et al., A brain-wide map of neural activity during complex behaviour. Nature, 2025. 645(8079): p. 177–191.

2. Nassar, M.R., et al., Age differences in learning emerge from an insufficient representation of uncertainty in older adults. Nature communications, 2016. 7(1): p. 11609.

3. Sevi, B., et al., Now You See Them, Now You Don’t: Age Differences in Risk Aversion. Innovation in Aging, 2020. 4(Suppl 1): p. 365.

4. Yang, T., C. Xie, and X. Wang, Forgetting phenomena in the Iowa Gambling Task: a new computational model among diverse participants. Frontiers in Psychology, 2025. **Volume** 16 **-** 2025.

5. Sun, T., et al., Decision-making under ambiguity or risk in individuals with Alzheimer’s disease and mild cognitive impairment. Frontiers in psychiatry, 2020. 11: p. 218.

6. Therrien, S., et al., Risk-taking behavior differs between older adults with and without mild cognitive impairment. Journal of Alzheimer’s Disease, 2024. 100(4): p. 1227–1235.

7. Bechara, A., et al., Insensitivity to future consequences following damage to human prefrontal cortex. Cognition, 1994. 50(1-3): p. 7–15.

8. Buelow, M.T. and J.A. Suhr, Construct validity of the Iowa Gambling Task. Neuropsychol Rev, 2009. 19(1): p. 102–14.

9. Lejuez, C.W., et al., Evaluation of a behavioral measure of risk taking: the Balloon Analogue Risk Task (BART). Journal of Experimental Psychology: Applied, 2002. 8(2): p. 75.

10. Wang, M., et al., Risk-taking in the human brain: An activation likelihood estimation meta-analysis of the balloon analog risk task (BART). Human brain mapping, 2022. 43(18): p. 5643–5657.

11. Compagne, C., et al., Tools for the assessment of risk-taking behavior in older adults with mild dementia: a cross-sectional clinical study. Brain sciences, 2023. 13(6): p. 967.

12. Skeel, R.L., et al., The utility of personality variables and behaviorally-based measures in the prediction of risk-taking behavior. Personality and Individual Differences, 2007. 43(1): p. 203–214.

13. Buelow, M.T. and W.R. Barnhart, The influence of math anxiety, math performance, worry, and test anxiety on the Iowa gambling task and balloon analogue risk task. Assessment, 2017. 24(1): p. 127–137.

14. Aklin, W.M., et al., Evaluation of behavioral measures of risk taking propensity with inner city adolescents. Behaviour research and therapy, 2005. 43(2): p. 215–228.

15. Lejuez, C., et al., The balloon analogue risk task (BART) differentiates smokers and nonsmokers. Experimental and clinical psychopharmacology, 2003. 11(1): p. 26.

16. Buelow, M.T. and A.L. Blaine, The assessment of risky decision making: a factor analysis of performance on the Iowa Gambling Task, Balloon Analogue Risk Task, and Columbia Card Task. Psychological assessment, 2015. 27(3): p. 777.

17. Frank, C.C. and K.L. Seaman, *Aging, uncertainty, and decision making—A review.* Cognitive, Affective, & Behavioral Neuroscience, 2023. 23(3): p. 773–787.

18. Hales, C.A., L. Clark, and C.A. Winstanley, Computational approaches to modeling gambling behaviour: Opportunities for understanding disordered gambling. Neuroscience & Biobehavioral Reviews, 2023. 147: p. 105083.

19. Worthy, D.A., B. Pang, and K.A. Byrne, Decomposing the roles of perseveration and expected value representation in models of the Iowa gambling task. Frontiers in Psychology, 2013. **Volume** 4 **-** 2013.

20. Fuchs, B.A., et al., Decision-Making Processes Related to Perseveration Are Indirectly Associated With Weight Status in Children Through Laboratory-Assessed Energy Intake. Frontiers in Psychology, 2021. **Volume** 12 **-** 2021.

21. Liu, Q., et al., Modulation of dlPFC function and decision-making capacity by repetitive transcranial magnetic stimulation in methamphetamine use disorder. Translational Psychiatry, 2024. 14(1): p. 280.

22. Molins, F., N. Ben Hassen, and M.Á. Serrano, Late acute stress effects on decision-making: The magnified attraction to immediate gains in the iowa gambling task. Behavioural Brain Research, 2025. 476: p. 115279.

23. Park, H., et al., Development of a novel computational model for the Balloon Analogue Risk Task: The exponential-weight mean–variance model. Journal of Mathematical Psychology, 2021. 102: p. 102532.

24. Qin, F., et al., Neural correlates of risk decision-making: Insights from the balloon analogue risk task and exponential-weight mean-variance model. Cortex, 2025. 187: p. 1–15.

25. Wittens, M.M.J., et al., Brain age as a biomarker for pathological versus healthy ageing – a REMEMBER study. Alzheimer’s Research & Therapy, 2024. 16(1): p. 128.

26. Kaufmann, T., et al., Common brain disorders are associated with heritable patterns of apparent aging of the brain. Nature Neuroscience, 2019. 22(10): p. 1617–1623.

27. Cox, S.R., et al., Brain cortical characteristics of lifetime cognitive ageing. Brain Structure and Function, 2018. 223(1): p. 509–518.

28. Li, W.-X., et al., White matter and gray matter changes related to cognition in community populations. Frontiers in Aging Neuroscience, 2023. **Volume** 15 **-** 2023.

29. Feng, S., et al., Brain networks under uncertainty: a coordinate-based meta-analysis of brain imaging studies. Journal of Affective Disorders, 2022. 319: p. 627–637.

30. Mf, F., A practical method for grading the cognitive state of patients for the clinician. J Psychiatr Res, 1992. 12: p. 189–198.

31. Nasreddine, Z.S., et al., The Montreal Cognitive Assessment, MoCA: a brief screening tool for mild cognitive impairment. Journal of the American Geriatrics Society, 2005. 53(4): p. 695–699.

32. Pinto, T.C., et al., Is the Montreal Cognitive Assessment (MoCA) screening superior to the Mini-Mental State Examination (MMSE) in the detection of mild cognitive impairment (MCI) and Alzheimer’s Disease (AD) in the elderly? International psychogeriatrics, 2019. 31(4): p. 491–504.

33. Cauffman, E., et al., Age differences in affective decision making as indexed by performance on the Iowa Gambling Task. Developmental psychology, 2010. 46(1): p. 193.

34. Ren, P., et al., Aging-related changes in reward-based decision-making depend on punishment frequency: An fMRI study. Frontiers in Aging Neuroscience, 2023. 15: p. 1078455.

35. Ren, P., et al., Dorsal and ventral fronto-amygdala networks underlie risky decision-making in age-related cognitive decline. GeroScience, 2024. 46(1): p. 447–462.

36. Ren, P., et al., Enhanced putamen functional connectivity underlies altered risky decision-making in age-related cognitive decline. Scientific reports, 2023. 13(1): p. 6619.

37. Haines, N., J. Vassileva, and W.Y. Ahn, The outcome-representation learning model: A novel reinforcement learning model of the iowa gambling task. Cognitive science, 2018. 42(8): p. 2534–2561.

38. Ahn, W.Y., et al., Comparison of decision learning models using the generalization criterion method. Cognitive science, 2008. 32(8): p. 1376–1402.

39. Ahn, W.-Y., et al., Decision-making in stimulant and opiate addicts in protracted abstinence: evidence from computational modeling with pure users. Frontiers in psychology, 2014. 5: p. 849.

40. Van Ravenzwaaij, D., G. Dutilh, and E.-J. Wagenmakers, Cognitive model decomposition of the BART: Assessment and application. Journal of mathematical psychology, 2011. 55(1): p. 94–105.

41. Yan, C.-G., et al., DPABI: Data Processing & Analysis for (Resting-State) Brain Imaging. Neuroinformatics, 2016. 14(3): p. 339–351.

42. Ashburner, J., A fast diffeomorphic image registration algorithm. Neuroimage, 2007. 38(1): p. 95–113.

43. Tzourio-Mazoyer, N., et al., Automated anatomical labeling of activations in SPM using a macroscopic anatomical parcellation of the MNI MRI single-subject brain. Neuroimage, 2002. 15(1): p. 273–289.

44. Yu, J., et al., Altered value coding in the ventromedial prefrontal cortex in healthy older adults. Frontiers in aging neuroscience, 2016. 8: p. 210.

45. Mata, R., et al., Age differences in risky choice: A meta-analysis. Annals of the new York Academy of Sciences, 2011. 1235(1): p. 18–29.

46. Albert, S.M. and J. Duffy, Differences in risk aversion between young and older adults. Neuroscience and neuroeconomics, 2012: p. 3–9.

47. Nagaratnam, J.M., et al., Trajectories of mini-mental state examination scores over the lifespan in general populations: a systematic review and meta-regression analysis. Clinical Gerontologist, 2022. 45(3): p. 467–476.

48. Coen, R.F., et al., Strengths and limitations of the MoCA for assessing cognitive functioning: findings from a large representative sample of Irish older adults. Journal of geriatric psychiatry and neurology, 2016. 29(1): p. 18–24.

49. Montagnese, M. and T. Rittman, Bridging modifiable risk factors and cognitive decline: the mediating role of brain age. Lancet Healthy Longev, 2024. 5(4): p. e243–e244.

50. Dias, M.F., et al., Unravelling pathological ageing with brain age gap estimation in Alzheimer’s disease, diabetes and schizophrenia. Brain Commun, 2025. 7(2): p. fcaf109.

51. Vassiliadis, P., et al., Non-invasive stimulation of the human striatum disrupts reinforcement learning of motor skills. Nat Hum Behav, 2024. 8(8): p. 1581–1598.

52. Lindsey, J.W., et al., Dynamics of striatal action selection and reinforcement learning. Elife, 2025. 13.

53. Neufeld, N., et al., Longitudinal changes in grey matter and cognitive performance over four years of healthy aging. Neuroimage Rep, 2022. 2(4): p. 100140.

54. Sawada, Y., et al., Assessment of localized brain regions correlated with MMSE using VBM analysis of structural MRI in a Japanese sample. Neuroimage Rep, 2025. 5(2): p. 100264.

55. Dörfel, R.P., et al., Prediction of brain age using structural magnetic resonance imaging: A comparison of accuracy and test–retest reliability of publicly available software packages. Human Brain Mapping, 2023. 44(17): p. 6139–6148.

