## Supplementary for "Estimating chronological and brain age using risk-taking behavior under uncertainty"

**Computational models in the IGT**

1. *Value-Plus-Perseveration (VPP) model*

The Value-Plus-Perseverance (VPP) model extends the basic reinforcement learning framework by incorporating a perseverance mechanism, which captures the tendency to repeat or avoid recently chosen options [1]. The model estimates eight parameters: Learning rate (0 < A < 1) determines the weight given to recent outcomes when updating expectancies; Feedback sensitivity (0 < alpha < 2) shapes the utility function for outcomes; Loss aversion (0 < lambda < 5) controls the subjective impact of losses relative to gains; Consistency (0 < cons < 5) reflects the determinism versus randomness of choices; Perseverance decay (0 < k < 1) reflects how the perseverance strength of all options in discounted over trials; Gain impact (epP) and loss impact (epN) modify the perseverance value after gains and losses, separately; Reinforcement-learning weight (0 < w < 1) balances the influence of reinforcement learning versus perseverance processes.

2. *Outcome-Representation Learning model (ORL)*

The ORL model characterizes decision-making by separately tracking the representations of gains and losses for each option [2]. It updates expected outcomes based on reinforcement learning principles while incorporating perseverance and exploration components. The model assumes that individuals maintain outcome-specific representations and adjust their choices according to prediction errors derived from experienced rewards and penalties. It includes following 5 parameters: Arew (reward learning rate), Apun (punishment learning rate), K (perseverance decay), betaF (outcome frequency weight), betaP (perseverance weight).

3. *Prospect Valence Learning model with the Delta rule (PVL-Delta)*

The PVL-Delta model integrates prospect theory with reinforcement learning to explain performance in the IGT[3]. In this model, outcomes are transformed into subjective utilities according to prospect theory, capturing differential sensitivity to gains and losses. Expected values for each deck are then updated using a delta learning rule based on prediction errors between experienced and expected outcomes. A softmax choice rule translates expected values into selection probabilities. It includes following 4 parameters: A (learning rate), alpha (outcome sensitivity), cons (response consistency), lambda (loss aversion).

4. *Prospect Valence Learning model with Decay (PVL-Decay)*

The PVL-Decay also combines prospect theory with reinforcement learning but differs in its learning mechanism [4]. Instead of updating only the chosen deck through prediction errors, this model assumes that the expected values of all decks gradually decay over time. After each trial, the experienced outcome modifies the chosen deck while unchosen decks lose value through decay. In this way, the PVL-Decay model captures how memory processes and recency effects influence decision-making during the Task. It includes following 4 parameters: A (decay rate), alpha (outcome sensitivity), cons (response consistency), lambda (loss aversion).

**Computational models in the BART**

1. *Exponential-Weight Mean-Variance (EWMV) Model*

For the BART, the Exponential-Weight Mean-Variance (EWMV) model formalizes risk-taking as a dynamic process where participants balance expected rewards against perceived risk [5]. The model assumes that participants continuously update their beliefs about the explosion probability and adjust their behavior accordingly. It is characterized by five parameters: Prior belief of burst (phi) represents the initial belief about the balloons explosion probability; Updating exponent (0 < eta < 1) acts as a learning rate for updating the belief about the explosion probability after each successful pump; Risk preference (rho) captures the individual preference for risk, influencing the weighting of expected value against variance; Inverse temperature (tau > 0) controls the randomness of decision-making with higher values indicating more deterministic choices. Loss aversion (lambda) modulates the subjective value of potential losses.

2. *4-parameter model*

The 4-parameter model assumes decision-makers update their belief in balloon burst probability using prior expectations and cumulative pump outcomes across trials [6]. It includes four latent parameters: phi (prior belief about explosion probability), representing the initial expectation about the balloon burst probability; eta (learning rate), reflecting how individuals adjust their belief about explosion risk based on outcomes across trials; gam (risk propensity), indicating the tendency to continue pumping despite potential loss; tau (behavioral consistency), describing the degree to which choices follow learned expectations versus random variability.

**References:**


1. Worthy, D.A., B. Pang, and K.A. Byrne, *Decomposing the roles of perseveration and expected value representation in models of the Iowa gambling task.* Frontiers in Psychology, 2013. **Volume 4 - 2013**.
2. Haines, N., J. Vassileva, and W.Y. Ahn, *The outcome-representation learning model: A novel reinforcement learning model of the iowa gambling task.* Cognitive science, 2018. **42**(8): p. 2534-2561.
3. Ahn, W.Y., et al., *Comparison of decision learning models using the generalization criterion method.* Cognitive science, 2008. **32**(8): p. 1376-1402.
4. Ahn, W.-Y., et al., *Decision-making in stimulant and opiate addicts in protracted abstinence: evidence from computational modeling with pure users.* Frontiers in psychology, 2014. **5**: p. 849.
5. Park, H., et al., *Development of a novel computational model for the Balloon Analogue Risk Task: The exponential-weight mean–variance model.* Journal of Mathematical Psychology, 2021. **102**: p. 102532.
6. Van Ravenzwaaij, D., G. Dutilh, and E.-J. Wagenmakers, *Cognitive model decomposition of the BART: Assessment and application.* Journal of mathematical psychology, 2011. **55**(1): p. 94-105.

**Table S1.** Correlation between computational parameters and chronological age in older adults.

| | IGT | | | | | | | | BART (high risk) | | | | | BART (low risk) | | | | |
|---|---|---|---|---|---|---|---|---|---|---|---|---|---|---|---|---|---|---|
| | A | alpha | cons | lambda | epP | epN | K | w | phi | eta | rho | tau | lambda | phi | eta | rho | tau | lambda |
| correlation | 0.075 | **-0.253** | -0.087 | 0.039 | **0.234** | -0.099 | **-0.264** | **0.198** | 0.056 | **0.217** | 0.028 | **-0.21** | 0.134 | 0.041 | 0.086 | 0.073 | -0.156 | 0.116 |
| *p value* | 0.431 | **0.007** | 0.362 | 0.680 | **0.013** | 0.299 | **0.005** | **0.037** | 0.555 | **0.022** | 0.773 | **0.026** | 0.158 | 0.665 | 0.369 | 0.446 | 0.101 | 0.222 |

**Table S2.** Correlation between computational parameters and brain age in older adults.

| | IGT | | | | | | | | BART (high risk) | | | | | BART (low risk) | | | | |
|---|---|---|---|---|---|---|---|---|---|---|---|---|---|---|---|---|---|---|
| | A | alpha | cons | lambda | epP | epN | K | w | phi | eta | rho | tau | lambda | phi | eta | rho | tau | lambda |
| correlation | 0.125 | **-0.251** | -0.044 | 0.075 | **0.213** | -0.084 | **-0.273** | 0.178 | 0.091 | **0.202** | -0.013 | -0.127 | 0.154 | 0.007 | 0.059 | 0.009 | -0.097 | 0.148 |
| *p value* | 0.188 | **0.008** | 0.642 | 0.429 | **0.024** | 0.379 | **0.004** | 0.060 | 0.339 | **0.033** | 0.893 | 0.183 | 0.106 | 0.941 | 0.535 | 0.926 | 0.309 | 0.118 |

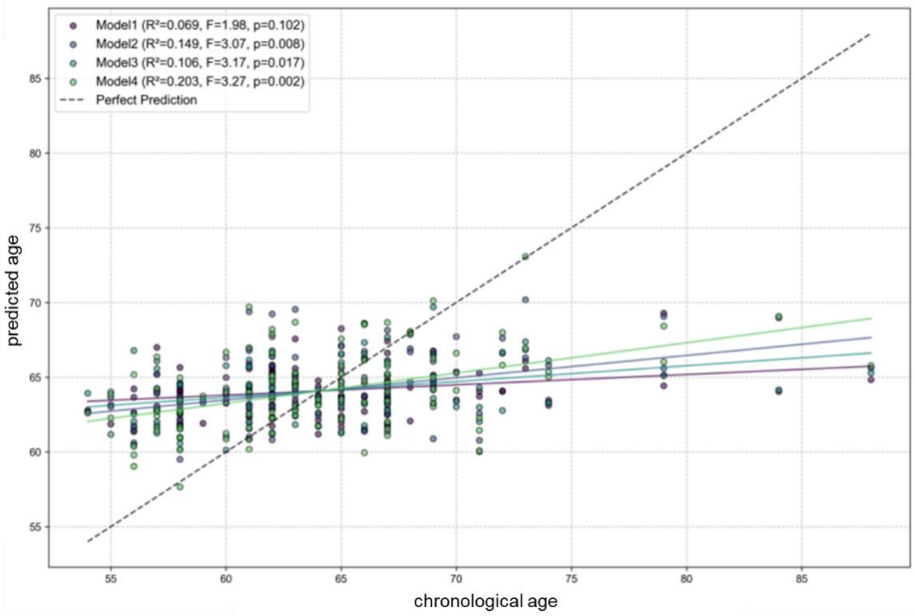


**Figure S1. Linear regression models in predicting chronological age.** For each model, $R^2$ value indicates the proportion of variance in actual age explained by the model, F value measures the overall significance of the model. Model 1: MMSE and MoCA; Model 2: VPP parameters in the IGT (i.e., alpha, epP, K, and w); Model 3: EWMV parameters in the BART (eta and tau in the high-risk condition); Model 4: model parameters in the IGT and BART (i.e., alpha, epP, K, w, $eta_{\text{high-risk}}$, and $tau_{\text{high-risk}}$).

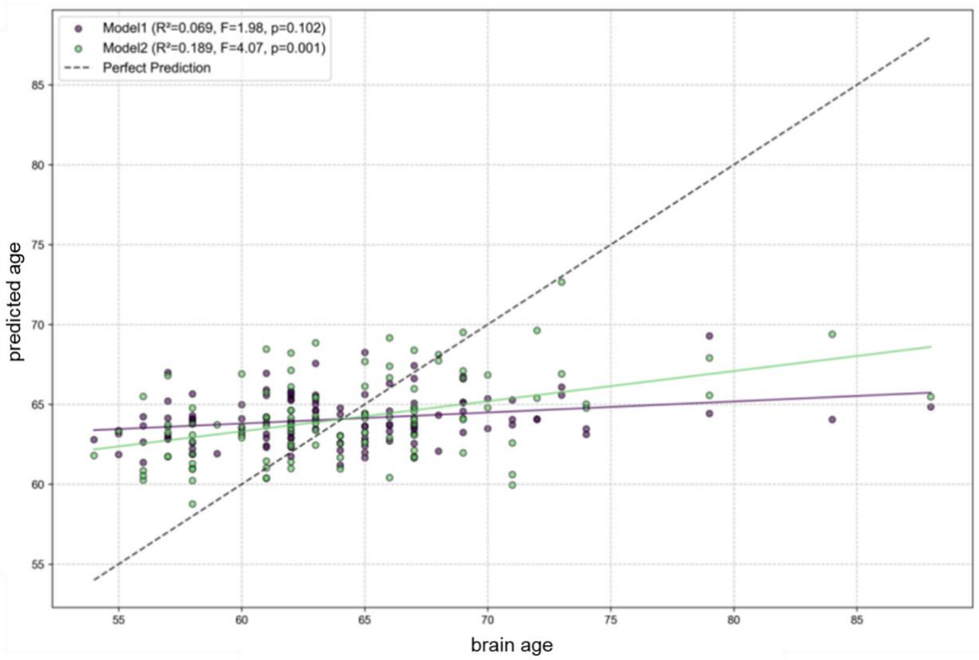


**Figure S2. Linear regression models in predicting brain age.** For each model, $R^2$ value indicates the proportion of variance in actual age explained by the model, F value measures the overall significance of the model. Model 1: MMSE and MoCA; Model 2: IGT + BART parameters (alpha, epP, K, eta $_{\text{high-risk}}$).

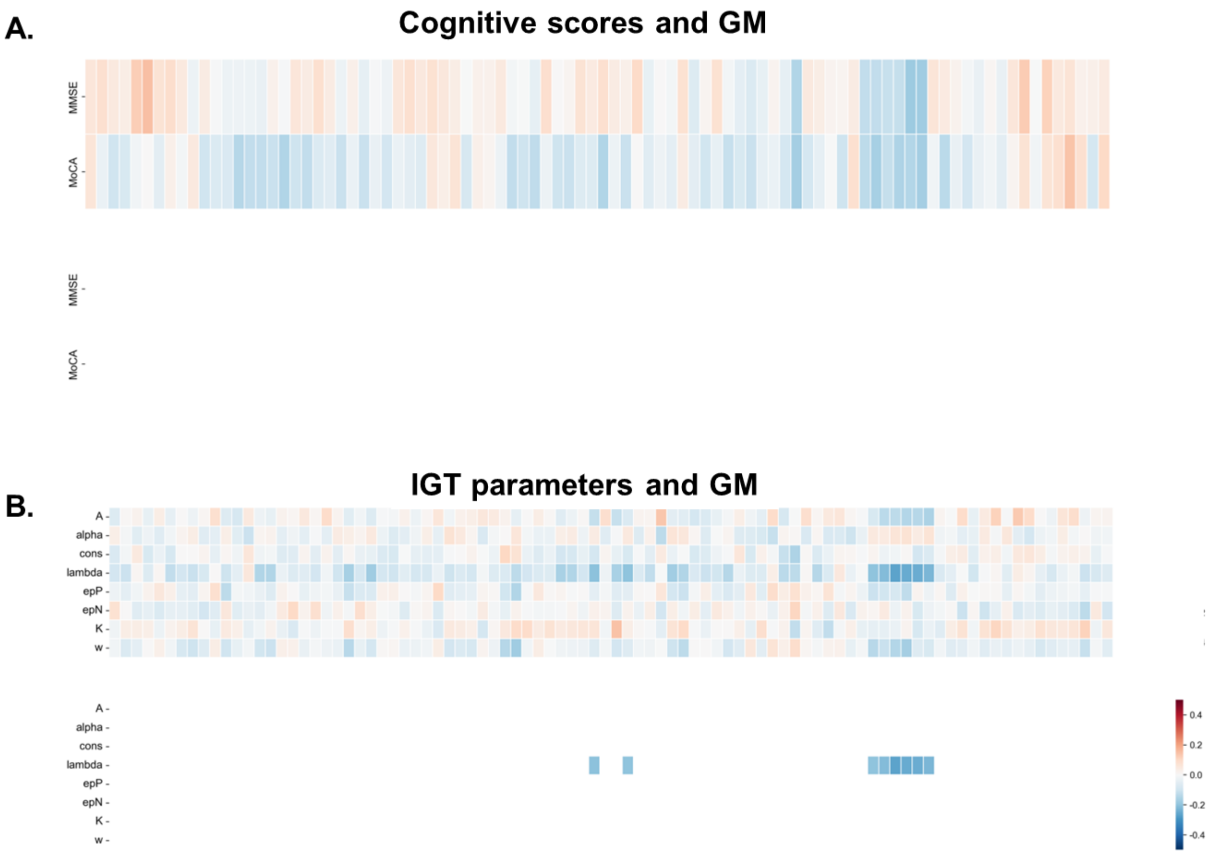


**Figure S3. Correlations between GM and cognitive scores/IGT parameters in older adults.** A) In MMSE/MoCA scores, no significant correlation was observed. B) In the IGT parameters, significant correlations were observed in the occipital and subcortical regions, which highlighted in the bottom matrix (uncorrected $p < 0.05$).

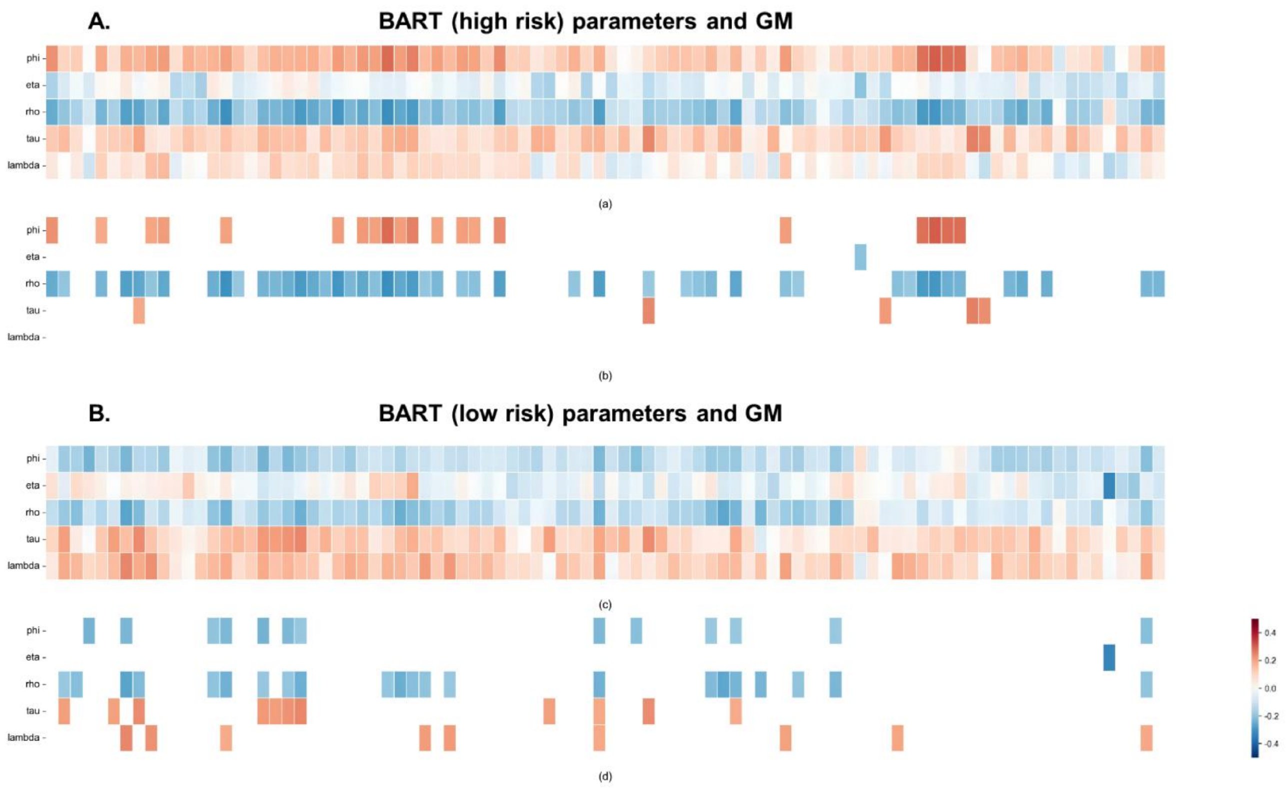


**Figure S4. Pearson correlations between BART parameters and GMV in older adults.** A) In the high risk condition, significant correlations were observed in multiple parameters, and significant brain regions were highlighted in the bottom matrix. B) In the low risk condition, significant correlations were observed in multiple parameters, and significant brain regions were highlighted in the bottom matrix. (uncorrected $p < 0.05$).
